# GLP-1 and GIP receptor agonism does not directly drive skeletal muscle atrophy or impair myogenesis in primary human myotubes

**DOI:** 10.64898/2026.07.20.739515

**Authors:** Caitlin Ditchfield, Michael Macleod, Joshua MJ Price, Edward T Davis, Simon W Jones

## Abstract

GLP-1 and GIP/GLP-1 receptor agonists produce substantial weight loss in clinical trials but significant loss of lean body mass is reported. Whether this reflects a direct pharmacological effect on skeletal muscle or an indirect consequence of caloric restriction and reduced mechanical loading is unknown.

Primary myoblasts were isolated from skeletal muscle of older adults with obesity undergoing orthopaedic surgery. GIPR and GLP-1R expression was characterised by RT-qPCR and flow cytometry. Differentiated myotubes were treated with semaglutide or GIP peptide and assessed for atrophy-related gene expression (qPCR), secretome perturbation (Olink Reveal), mitochondrial and glycolytic bioenergetics (Seahorse XF Real-Time ATP Rate Assay, glucose uptake, lactate secretion) and myotube morphology and myogenesis (immunofluorescence).

GIPR mRNA was consistently detected across all donors; GLP-1R mRNA was undetectable by PCR, though LUXendin645 flow cytometry identified low-level surface GLP-1R protein in 51–66% of myoblasts. Neither semaglutide nor GIP altered atrophy-related gene expression or the secretome, with no proteins reaching significance. Semaglutide reduced glycolytic and total ATP production rates, accompanied by reduced lactate secretion, suggesting modest suppression of glycolytic flux; mitochondrial parameters were unaffected. Neither treatment impaired myotube thickness or differentiation; GIP increased myotube thickness after 8 days.

Direct GLP-1 and GIP receptor activation does not substantively perturb atrophic signalling, myogenesis, or the secretome of primary human skeletal muscle myotubes. These findings suggest that lean mass loss with incretin-based therapies is unlikely to be driven by direct pharmacological action on skeletal muscle - particularly relevant as these agents are increasingly used in older adults at risk of sarcopenia.

## Background

Glucagon-like peptide-1 (GLP-1) receptor agonists including semaglutide and liraglutide and the dual GLP-1/glucose-dependent insulinotropic polypeptide (GIP) receptor agonist tirzepatide have transformed the clinical management of type 2 diabetes mellitus (T2DM) and obesity. Acting on incretin receptors, they have been shown to potentiate glucose-stimulated insulin secretion, suppress glucagon release, delay gastric emptying, and reduce appetite, collectively resulting in substantial reductions in body weight and improved glycaemic control. Importantly, clinical trials including SUSTAIN (1), STEP (2), LEADER (3), SELECT (4), and SURMOUNT (5) have demonstrated benefits not only in reducing body weight and improving glycaemic control but also reductions in adverse cardiovascular events including heart failure, and all-cause mortality.

Importantly, despite their well-established benefits, clinical trial data has shown that these agents also result in a reduction in lean body mass i.e., skeletal muscle mass (6). For example, trials with semaglutide in over-weight and obese adults have shown up to a 9% reduction in lean mass (7), whilst an approximately 11% decline in lean mass was found with patients on tirzepatide (5). Although absolute fat mass declined to a greater extent, and as such fat to lean ratio was improved, the absolute loss of lean mass has raised concerns given the critical functional role of skeletal muscle in physical function/mobility and in metabolic homeostasis. These concerns are especially pertinent in the context of older adults due to the progressive, generalised loss of skeletal muscle mass and function with age, termed sarcopenia. Sarcopenia affects an estimated 10–20% of adults over 60 years and is associated with increased risk of falls, frailty, disability, hospitalisation, and premature mortality (8).

Crucially, with an increasingly aged and obese population, the coexistence of excess adiposity and low muscle mass/function with age, termed sarcopenic obesity, is increasingly prevalent and confers a compounded metabolic and functional risk beyond either condition alone (9). A recent meta-analysis found that 11% of individuals >65 years old are comorbid with sarcopenia and obesity (10), and exhibit increased risk of cardiovascular disease, osteoporosis-linked fractures, Type 2 Diabetes, hospitalisation and all-cause mortality (11–13). In this context, therapies that produce substantial weight loss but also erode muscle mass could, in susceptible individuals, accelerate functional decline even while improving cardiometabolic parameters.

At present, the mechanism by which incretin agonists reduce lean mass remains poorly understood. One possibility is that lean mass loss is secondary to the caloric deficit and reduced mechanical loading accompanying overall weight reduction, rather than a direct pharmacological adverse effect on skeletal muscle. However, this has not been conclusively established, and the possibility of direct incretin receptor-mediated effects on muscle biology warrants investigation. GLP-1 receptors (GLP-1R) have been reported in rodent skeletal muscle (14), and preclinical studies have suggested that GLP-1 agonists may influence skeletal muscle glucose uptake, muscle protein synthesis, and inflammatory signalling (14). With regards to GIP, expression of the GIP receptor (GIPR) has not been consistently identified in whole skeletal muscle or liver. However, GIP agonists have been shown to stimulate glucose transport in rodent muscle independently of classical GIPR signalling, and GIPR mRNA has been detected in fibro-adipogenic progenitor cells within mouse muscle via in situ hybridisation (15–17). Therefore, the aim of the present study was firstly to evaluate the expression of GLP-1R and GIPR in human muscle tissues from older over-weight and obese adults and secondly to comprehensively determine whether GLP-1 and GIP receptor agonists exert direct effects on primary human skeletal muscle myotube atrophic signalling, growth and differentiation and metabolic function.

## Methods

### Patient Recruitment and Sample Collection

Adipose and skeletal muscle tissues were obtained from over-weight/obese (mean BMI 32.48 ± 2.58) older adults (mean age 73 ± 7.17) undergoing elective total knee or hip arthroplasty for osteoarthritis at the Royal Orthopaedic Hospital (Birmingham, UK) and Russell’s Hall Hospital (Dudley, UK), following ethical approval (NRES 17/SS/0456; 16/SS/0172), and written informed consent from all participants. Clinical metadata including age, BMI, and current medications were recorded. None of the participants were receiving incretin analogues or DPP-4 inhibitors prior to sample collection.

### Isolation, Culture, and Differentiation of Primary Human Myoblasts

Fresh muscle tissue was finely minced using sterile scalpels and digested in 5 mL 1x trypsin for 15 min at 37°C. Digestion was quenched by adding an equal volume of myoblast growth medium (Ham’s F-10; Sigma N6013, supplemented with 2 mM L-glutamine, 100 µg/mL penicillin/streptomycin, and 20% heat-inactivated FBS). The suspension was centrifuged at 520 x g for 5 min and the resulting pellet resuspended in growth medium. To deplete fibroblasts, the suspension was pre-plated on uncoated flasks for 20 min at 37°C. The non-adherent fraction was transferred onto 0.2% gelatin-coated flasks or plates (gelatin Type B 2% stock; Sigma G1393, diluted 1:10 in PBS). Cells were maintained at 37°C, 5% CO_₂_, with medium replaced every 2–3 days and passaged at 80% confluence. Early-passage cells (≤P3) were used for all experiments. At confluence, growth medium was replaced with differentiation medium (Ham’s F-10 supplemented with 6% horse serum). Medium was changed every 48 h for 8 days. Myogenic differentiation was confirmed by the appearance of elongated, multinucleated myotubes.

### Cell Treatments

Differentiated human skeletal muscle myotubes (n = 6 donors) were treated with semaglutide (10 nM; MedChemExpress HY-114118; DMSO vehicle) or GIP peptide (5.02 µM; MedChemExpress HY-P0276; H_₂_O vehicle). GIP and H_₂_O vehicle control conditions additionally received the DPP-4 inhibitor DPP4-010 (Sigma-Aldrich) at a final concentration of 50 µM to prevent GIP degradation.

### RNA Isolation and Quantitative PCR

RNA was extracted using the RNeasy Mini Kit (Qiagen). Concentration and purity were assessed using a NanoDrop One (Thermo Fisher Scientific, UK), and only samples with A260/280 ratios of 1.8–2.0 were included. For TaqMan-based targets (GIPR, MAFBX, MURF1, FOXO3, IL6, TGFB, GLP1R), qPCR was performed using the TaqMan One-Step RT-qPCR Kit (Bio-Rad) with gene-specific TaqMan probes (assay IDs listed in Supplementary Table S1). For SYBR-based targets (MYOG, MYOD, MYF5), reactions were performed using SYBR Green chemistry with gene-specific primers (sequences listed in Supplementary Table S1). All reactions contained 5 ng RNA in a 5 µL final volume and were run in triplicate on a CFX Opus Real-Time PCR System (Bio-Rad). GAPDH was used as the reference gene and relative expression calculated by 2-ΔΔCt.

### GLP-1R Flow Cytometry

Human primary myoblasts were incubated with LUXendin645 (500 nM) (18) for 15 min at room temperature to label surface GLP-1R. Cells were analysed on a BD Accuri C6 Plus flow cytometer. Gating was performed sequentially: debris exclusion by FSC/SSC scatter, singlet selection, and identification of LUXendin645-positive cells by fluorescence. Data are expressed as mean fluorescence intensity (MFI) fold change relative to unstained controls and percentage LUXendin645-positive cells.

### Olink Proteomics

Conditioned media were collected after 24 hours of treatment with semaglutide (DMSO vehicle) or GIP (H_₂_O vehicle), clarified by centrifugation at 13,000 × g for 2 min, aliquoted, and stored at −80°C prior to shipment to Birmingham Genomics Core. Samples were analysed using the Olink Reveal proximity extension assay panel (consisting of 1,034 proteins) according to standard workflows. Normalised protein expression (NPX) values, quality-controlled by the provider, were returned for 24 samples (n = 6 donors per condition: GIP, H_₂_O vehicle control, semaglutide, DMSO vehicle control). Samples with AssayQC = PASS were retained.

For visualisation, NPX values were scaled and principal component analysis (PCA) was performed in Python using scikit-learn (v1.0). Differential protein expression between each treatment and its respective vehicle control was assessed by two-sided Welch’s independent-samples t-test across all 1,034 proteins. Multiple testing correction was applied using the Benjamini–Hochberg procedure. Volcano plots were generated using log_₂_ fold change (calculated as the difference in mean NPX between groups, as NPX values are log_₂_- scaled) against −log_₁₀_ nominal p-value. Heatmaps display z-scored NPX values for the 100 proteins with the lowest nominal p-values per comparison, with rows hierarchically clustered using Ward linkage on Euclidean distances. Delta NPX values for selected proteins of interest were calculated as NPX(sample) − mean NPX(vehicle control) and plotted as individual values with mean ± SEM. All visualisations were produced in Python (matplotlib, seaborn) or R (ggplot2, pheatmap).

### Quantification of myogenesis and myotube thickness

Myotubes were differentiated in 24-well plates and then treated for 24 h with either semaglutide or GIP with appropriate vehicle controls. Cells were fixed in 2% paraformaldehyde for 30 min, washed, and stored in PBS. Cells were permeabilised with 100% methanol for 10 min, blocked with 5% goat serum (Gibco 16210064) for 30 min, and stained overnight at 4°C with rabbit anti-desmin (Abcam ab15200; RRID AB_301744; 1:1000 in 1% BSA/PBS). FITC-conjugated goat anti-rabbit IgG (Thermo Fisher Scientific F-2765; RRID AB_2536525; 1:1000) was applied for 1 h at room temperature in the dark, followed by DAPI (1:5000; Abcam ab228549) for 5 min. PBS was added to samples for imaging. Images were acquired on a Leica DMI6000 epifluorescence microscope (×20 and ×63 objective; 10 fields per well). Myotube thickness was quantified by averaging five transverse diameter measurements per myotube. Nuclei fusion index (NFI) was defined as the percentage of nuclei contained within desmin-positive multinucleated cells (≥3 nuclei). Myotube coverage was calculated as the percentage of total image area occupied by desmin-positive myotubes. All measurements were performed in ImageJ and using MyoCount (19) on 10 fields per well.

### Myotube metabolic functional assays

Real-time cellular energy metabolism of myotubes was quantified using the Seahorse XF Analyzer. Myotubes were differentiated in Seahorse XF96 96-well plates and treated with semaglutide or GIP with appropriate vehicle controls for 24 h. Mitochondrial and glycolytic ATP production rates were measured using the Seahorse XF Real-Time ATP Rate Assay Kit according to the manufacturer’s instructions. Data were normalised to total protein content (µg protein per well). Oxygen consumption rate (OCR), glycolytic proton efflux rate (GlycoPER), mitochondrial ATP production rate, glycolytic ATP production rate, total ATP production rate, coupling efficiency, proton leak, and non-mitochondrial respiration were calculated using Seahorse Wave software.

Glucose uptake was determined using the 2NBDG glucose uptake assay. Myotubes were treated with semaglutide or GIP with appropriate vehicle controls for 24 h; cells were then washed and incubated in glucose free media for 3 hr. 2NBDG (Thermo Fisher) was spiked in at 100 µM for the final 30 min of the 3 h glucose-starvation at 37°C. Fluorescence was measured on a plate reader (excitation 485 nm, emission 528 nm) and values normalised to total protein content.

L-Lactate concentrations in conditioned media were measured using the MAK570 L-Lactate Assay Kit (Sigma-Aldrich) according to the manufacturer’s instructions. Myotubes were treated for 24 h, washed with PBS and fresh media was added for 3 h. Values were normalised to total protein content and expressed as ng/mL.

### Data and Statistical Analysis

Data are presented as mean ± SD unless otherwise stated. For experiments involving multiple donors, comparisons between treatment and vehicle control were made by paired t-test or Wilcoxon signed-rank test as appropriate. For experiments from a single donor with biological replicates, unpaired comparisons were made by Wilcoxon signed-rank test. Statistical analyses were performed in GraphPad Prism v10. A p-value of < 0.05 was considered statistically significant.

## Results

### Primary human myotubes express GIPR and GLP-1R

To determine whether human skeletal muscle myotubes express incretin receptors, we assessed GIPR and GLP-1R mRNA expression by RT-PCR in differentiated primary myotubes from six older adults with BMI >30 (obese, Figure 1A). GIPR was consistently detected across all donors, with ΔCT values ranging from 33–36 cycles (mean ± SD per donor: 35.00 ± 0.81, 33.07 ± 0.71, 35.01 ± 0.73, 34.85 ± 1.21, 35.80 ± 0.54, 33.36 ± 0.23; Figure 1B). In contrast, GLP-1R mRNA was undetectable in all donors under the same assay conditions (Figure 1B).

**Figure 1.**
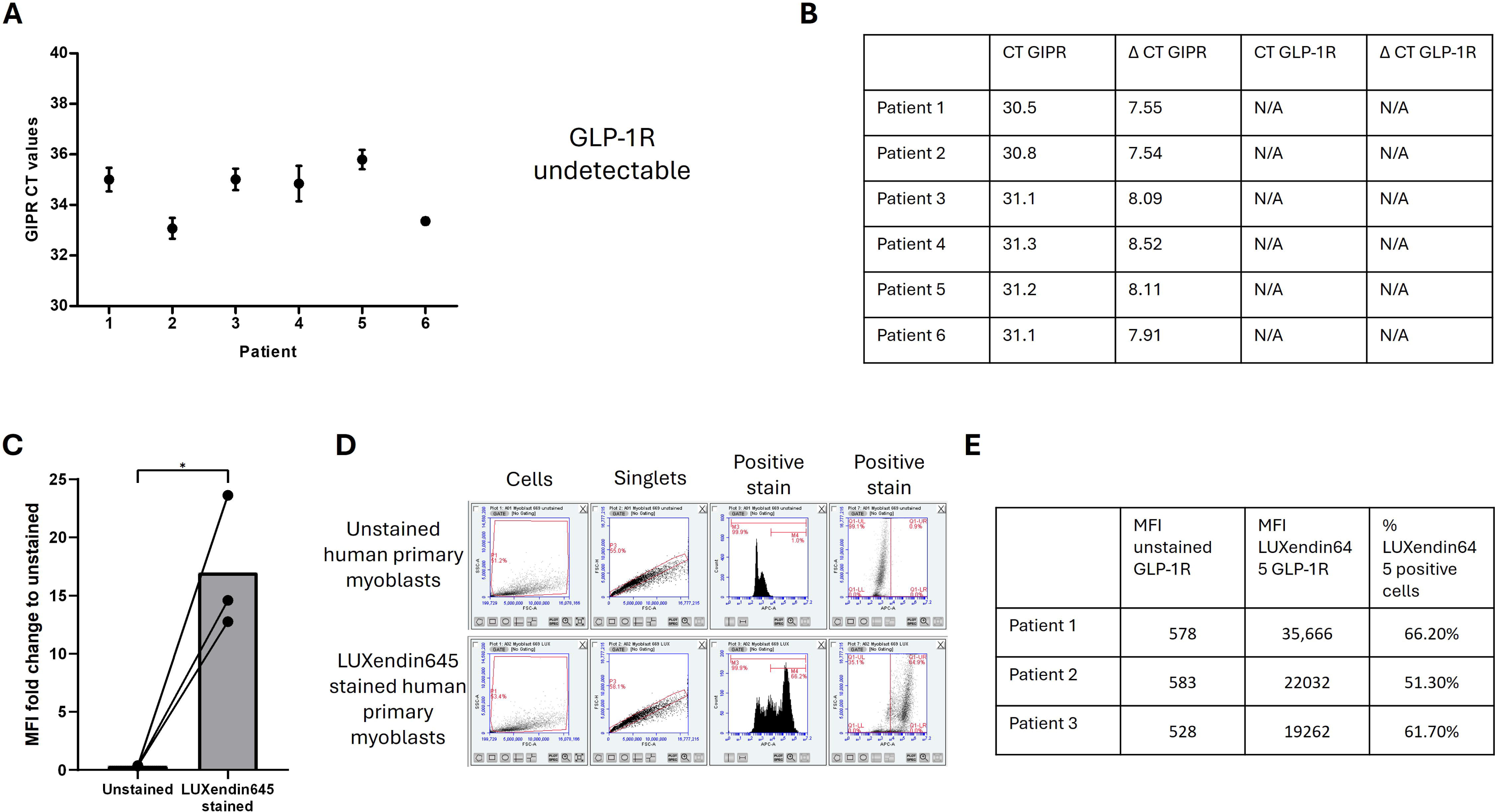
Primary human myotubes express GIPR but not GLP-1R. (A) RT-PCR analysis of GIPR and GLP-1R expression in differentiated primary human myotubes from six osteoarthritis patients (three technical replicates per donor). GIPR was consistently detected across all donors; GLP-1R was undetectable in all samples. (B) CT and ΔCT values for GIPR and GLP-1R across all six donors (ΔCT values: 35.00 ± 0.81, 33.07 ± 0.71, 35.01 ± 0.73, 34.85 ± 1.21, 35.80 ± 0.54, 33.36 ± 0.23; mean ± SD). (C) Mean fluorescence intensity (MFI) fold change relative to unstained controls in human primary myoblasts live-stained with LUXendin645 (500 nM, 15 minutes). Data from three donors shown with individual patient values connected by lines; bar represents the group mean. *p = 0.0383, paired t-test. (D) Representative gating strategy for LUXendin645 flow cytometry on the BD Accuri C6 Plus. Upper row: unstained human primary myoblasts. Lower row: LUXendin645-stained human primary myoblasts. Sequential gating: (1) FSC/SSC scatter to exclude debris, (2) singlet discrimination, (3 & 4) fluorescent signal as histogram and dot plot to identify LUXendin645-positive cells. (E) MFI values and percentage LUXendin645-positive cells for each of the three donors.

To investigate whether GLP-1R is expressed in human primary myotubes at the protein level, we performed live-cell flow cytometry using LUXendin645, a validated fluorescent GLP-1R antagonist probe (20). Despite the absence of GLP-1R mRNA by PCR, LUXendin645 staining produced a significant increase in MFI compared with unstained controls across three donors (MFI fold change: 16.98 ± 5.81; p = 0.0383, paired t-test; Figure 1C) with between 51–66% of cells classified as LUXendin645-positive. These data suggest that human primary myoblasts express low levels of surface GLP-1R protein that fall below the detection threshold of PCR. Representative gating strategy is shown in Figure 1D and individual MFI values in Figure 1E.

### Direct incretin treatment does not cause atrophic or inflammatory signalling dysregulation in muscle

To assess whether direct incretin exposure alters early transcriptional markers of atrophy or inflammation, human primary myotubes from six donors were treated with semaglutide or GIP for 4 hours and mRNA expression of GIPR, MAFBX, MURF1, FOXO3, IL6, and TGFB was quantified by qPCR. GLP-1R was also assessed but remained undetectable in all samples, consistent with Figure 1A. No significant differences were observed for any gene in either the semaglutide or GIP treatment conditions compared with their respective vehicle controls (Figure 2A). These data indicate that direct incretin exposure does not induce early transcriptional changes in canonical atrophy- or inflammation-related pathways in human myotubes.

**Figure 2.**
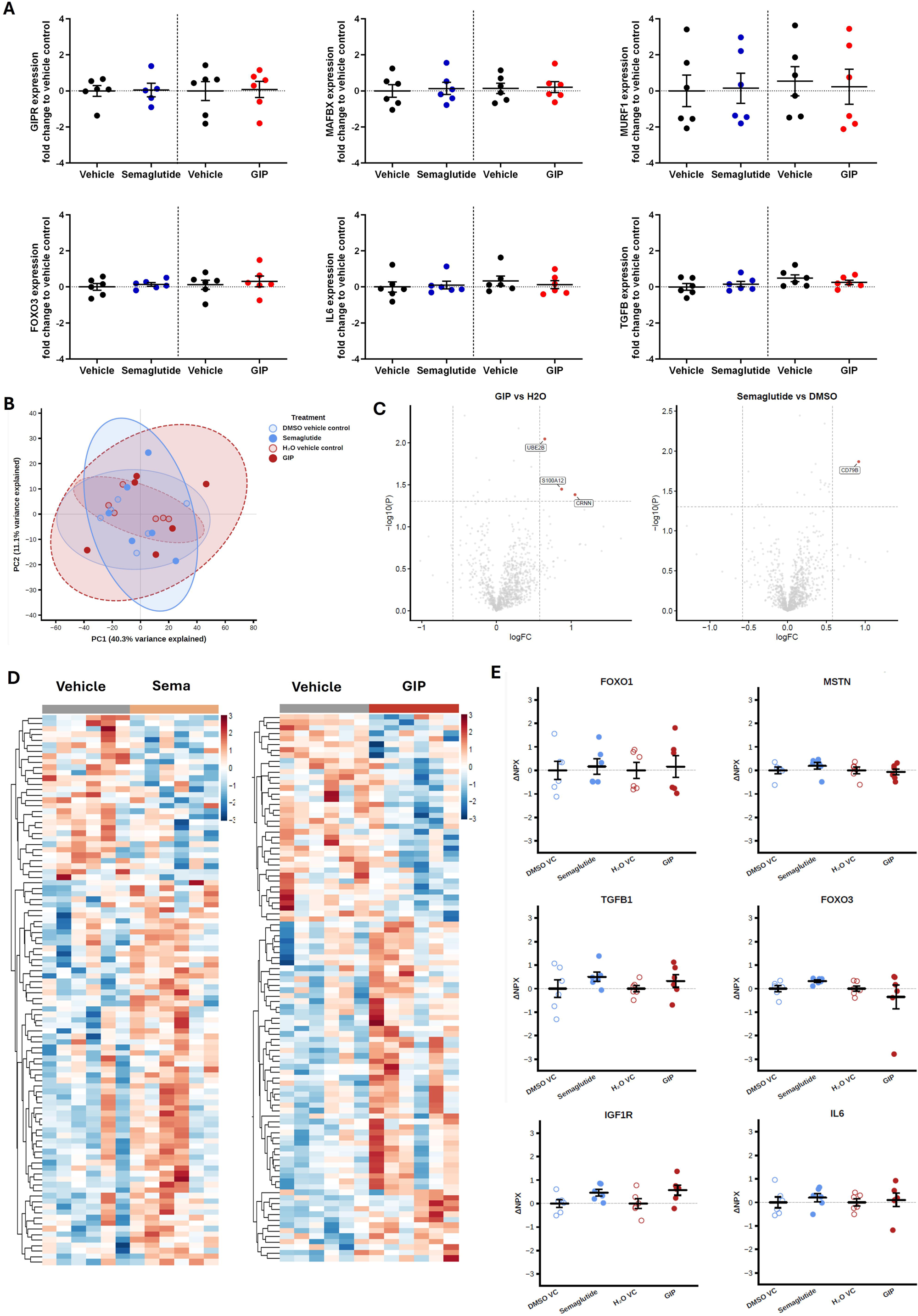
Incretins do not directly cause atrophic or inflammatory signalling dysregulation in muscle. Data from n = 6 donors unless otherwise stated. All PCR comparisons by Wilcoxon signed-rank test; ns = not significant. (A) mRNA expression (fold change to vehicle control) of atrophy- and inflammation-related genes in human primary myotubes 4 hours after treatment with semaglutide vs DMSO (left) or GIP vs H_₂_O (right). Genes assessed: GIPR, MAFBX, MURF1, FOXO3, IL6, and TGFB. GLP-1R was also assessed but was undetectable in all samples, consistent with Figure 1A. (B) Principal component analysis (PCA) of Olink Reveal normalised protein expression (NPX) data from primary human myotubes treated with GIP (H_₂_O vehicle control), semaglutide (DMSO vehicle control), or their respective vehicle controls. Points represent individual samples; 95% confidence ellipses are shown for each treatment group. (C) Volcano plots showing differential protein expression between GIP vs H_₂_O vehicle control (left) and semaglutide vs DMSO vehicle control (right). The x-axis shows log_₂_ fold change (NPX) and the y-axis shows −log_₁₀_ (nominal p-value) from Welch’s two-sample t-test. Dashed lines indicate thresholds of |log_₂_FC| > 0.58 and p < 0.05 (unadjusted). No proteins reached statistical significance after Benjamini–Hochberg correction across 1,034 proteins tested (minimum BH-adjusted p = 1.0, both comparisons). At nominal thresholds, three proteins were upregulated in GIP-treated myotubes (UBE2B, S100A12, CRNN), and one following semaglutide treatment (CD79B). (D) Heatmaps showing z-scored NPX values for the 100 proteins with the lowest nominal p-values from GIP vs H_₂_O comparison (left) and semaglutide vs DMSO comparison (right). Rows are hierarchically clustered; columns ordered by treatment group. Colour scale: red = high NPX, blue = low NPX. (E) Individual delta NPX values (NPX − mean NPX of vehicle control) for six proteins of functional relevance to muscle biology: FOXO1, MSTN, IGF1R, TGFB1, FOXO3, and IL6. Data shown for semaglutide vs DMSO vehicle control and GIP vs H_₂_O vehicle control. Individual donor values are plotted with mean ± SEM.

To obtain an unbiased assessment of the myotube secretome following incretin treatment, conditioned media were analysed using the Olink Reveal proximity extension assay platform, providing quantitative NPX data for 1,034 proteins (Supplementary File 1 and 2). Quality control density plots confirmed consistent data quality across all samples (Supplementary figure 1). Principal component analysis of scaled NPX data showed that samples did not segregate by treatment condition: PC1 explained 40.3% and PC2 explained 11.1% of total variance, with the 95% confidence ellipses of all four groups overlapping extensively (Figure 2B). This indicates that neither semaglutide nor GIP treatment produces a dominant shift in the myotube secretory proteome relative to vehicle control.

Differential expression analysis by Welch’s t-test, followed by Benjamini–Hochberg correction, identified no proteins reaching statistical significance at an adjusted p-value threshold of 0.05 in either comparison (minimum BH-adjusted p = 1.0 for both GIP vs H_₂_O and semaglutide vs DMSO; Figure 2C). At the level of nominal p-value (p < 0.05) combined with a |log_₂_FC| threshold of 0.58 (equivalent to a ∼1.5-fold change in NPX), three proteins were nominally upregulated following GIP treatment: UBE2B (log_₂_FC = 0.65, p = 0.009), S100A12 (log_₂_FC = 0.87, p = 0.036), and CRNN (log_₂_FC = 1.05, p = 0.042, Figure 2C). For semaglutide, one protein met both thresholds: CD79B (log_₂_FC = 0.92, p = 0.014, Figure 2C). Heatmaps of the top 100 proteins by nominal p-value in each comparison further illustrate that differential signals are modest and interspersed with substantial inter-donor variability (Figure 2D).

For six proteins with established relevance to muscle biology - FOXO1, MSTN, IGF1R, TGFB1, FOXO3, and IL6 - delta NPX values (relative to mean vehicle control) were plotted for all donors (Figure 2E). None of these comparisons reached statistical significance: FOXO1 (GIP: log_₂_FC = 0.16, p = 0.783; semaglutide: log_₂_FC = 0.16, p = 0.753), MSTN (GIP: log_₂_FC = −0.07, p = 0.739; semaglutide: log_₂_FC = 0.19, p = 0.352), IGF1R (GIP: log_₂_FC = 0.57, p = 0.086; semaglutide: log_₂_FC = 0.46, p = 0.057), TGFB1 (GIP: log_₂_FC = 0.32, p = 0.318; semaglutide: log_₂_FC = 0.50, p = 0.278), FOXO3 (GIP: log_₂_FC = −0.35, p = 0.526; semaglutide: log_₂_FC = 0.32, p = 0.060), and IL6 (GIP: log_₂_FC = 0.10, p = 0.758; semaglutide: log_₂_FC = 0.20, p = 0.505). Together, these data indicate that direct activation of incretin pathways does not significantly alter the secretion of canonical muscle atrophy, growth, or inflammation markers under basal conditions.

### Direct incretin treatment has divergent effects on skeletal muscle glycolytic activity and metabolic substrate utilisation

To assess whether incretins directly alter skeletal muscle energy metabolism, mitochondrial and glycolytic bioenergetics were measured using the Seahorse XF Real-Time ATP Rate Assay in myotubes from four donors treated with semaglutide or GIP peptide. Neither treatment affected oxygen consumption rate (OCR; semaglutide p = 0.2764, GIP p = 0.1295; Figure 3A) or basal mitochondrial ATP production rate (J_ATP-MITO; semaglutide p = 0.3160, GIP p = 0.4797; Figure 3B), indicating that oxidative phosphorylation was unaffected.

**Figure 3.**
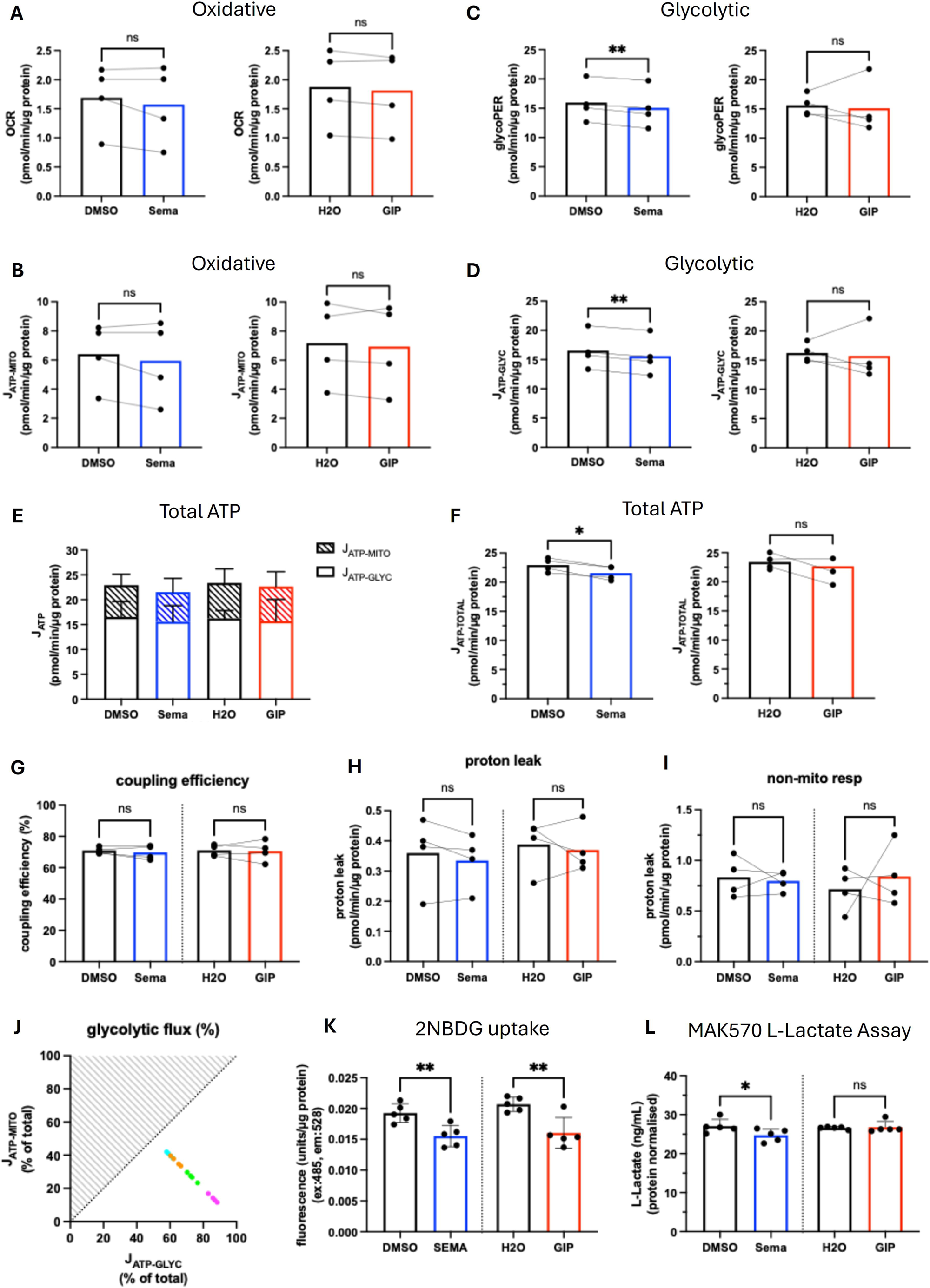
Incretins do not directly influence skeletal muscle cellular bioenergetics or metabolism. Panels A–J: n = 4 donors, paired t-test. Panels K–L: n = 5 biological replicates from a single donor, Wilcoxon signed-rank test. ns = not significant. (A) Oxygen consumption rate (OCR) for semaglutide vs DMSO (left) and GIP vs H_₂_O (right). (B) (B) Mitochondrial ATP production rate (J_ATP-MITO) for semaglutide vs DMSO (left) and GIP vs H_₂_O (right). (C) (C) Glycolytic proton efflux rate (GlycoPER) for semaglutide vs DMSO (left) and GIP vs H_₂_O (right). Semaglutide p = 0.0035; GIP p = 0.7734. (D) Glycolytic ATP production rate (J_ATP-GLYC) for semaglutide vs DMSO (left) and GIP vs H_₂_O (right). Semaglutide p = 0.0014; GIP p = 0.7634. (E) Total ATP production rate showing mitochondrial (hatched) and glycolytic (open) contributions across all four conditions. (F) Total ATP production rate for semaglutide vs DMSO (left) and GIP vs H_₂_O (right). Semaglutide p = 0.0187; GIP p = 0.6688. (G) Mitochondrial coupling efficiency (%) across all four conditions. (H) Proton leak (pmol/min/µg protein) across all four conditions. (I) Non-mitochondrial respiration (pmol/min/µg protein) across all four conditions. (J) Bioenergetic space plots showing mitochondrial vs glycolytic ATP production as percentage of total (left) and absolute values (right). Each colour represents an individual donor, illustrating that inter-donor variation exceeds treatment effect. (K) 2NBDG fluorescent glucose uptake (ex:485, em:528), protein-normalised, measured 3 hours post-treatment with semaglutide vs DMSO (left) and GIP vs H_₂_O (right). Semaglutide p = 0.0095; GIP p = 0.0017. (L) L-Lactate concentration (ng/mL, protein-normalised) measured 3 hours post-treatment with semaglutide vs DMSO (left) and GIP vs H_₂_O (right). Semaglutide p = 0.0414; GIP p = 0.9724.

In contrast, semaglutide significantly reduced both glycolytic proton efflux rate (GlycoPER; p = 0.0035; Figure 3C) and glycolytic ATP production rate (J_ATP-GLYC; p = 0.0014; Figure 3D), whilst GIP had no effect on either measure (GlycoPER p = 0.7734; J_ATP-GLYC p = 0.7634). Total ATP production, comprising mitochondrial and glycolytic contributions, is shown for all four conditions (Figure 3E). Consistent with this, total ATP production rate was significantly reduced by semaglutide (p = 0.0187; Figure 3F) but not by GIP (p = 0.6688; Figure 3F).

Mitochondrial coupling efficiency (semaglutide p = 0.7134, GIP p = 0.9840), proton leak (semaglutide p = 0.4658, GIP p = 0.8961), and non-mitochondrial respiration (semaglutide p = 0.9354, GIP p = 0.8642) were unaffected by either treatment (Figures 3G–I). Bioenergetic space plots revealed that inter-donor variability was a greater source of variability than treatment effect (Figure 3J).

To complement the Seahorse data, glucose uptake and lactate production were assessed in a single donor using five biological replicates. 2NBDG fluorescent glucose uptake was measured at 3 hours post-treatment. Both semaglutide (p = 0.0095) and GIP (p = 0.0017) significantly reduced glucose uptake relative to their respective vehicle controls (Figure 3K). L-Lactate concentrations in conditioned media were significantly reduced by semaglutide (p = 0.0414) but were unaffected by GIP (p = 0.9724; Figure 3L).

Together, these findings indicate that semaglutide suppresses glycolytic flux and reduces glucose uptake and lactate production in skeletal muscle myotubes, whereas GIP reduces glucose uptake without measurably altering downstream glycolytic or bioenergetic outputs. The mechanistic basis and functional significance of these divergent effects warrant further investigation.

### Direct incretin treatment does not cause myotube atrophy

To confirm the validity of our atrophy model, differentiated myotubes were stained with desmin and DAPI. Undifferentiated myoblasts appeared as long, thin, mononucleated cells, whereas differentiated myotubes exhibited the expected multinucleated, elongated morphology (Figure 4A). Treatment with TNFα for 24 hours induced visible myotube thinning whilst preserving multinucleation, confirming that the system responds appropriately to a well-established atrophic stimulus (Figure 4B).

**Figure 4.**
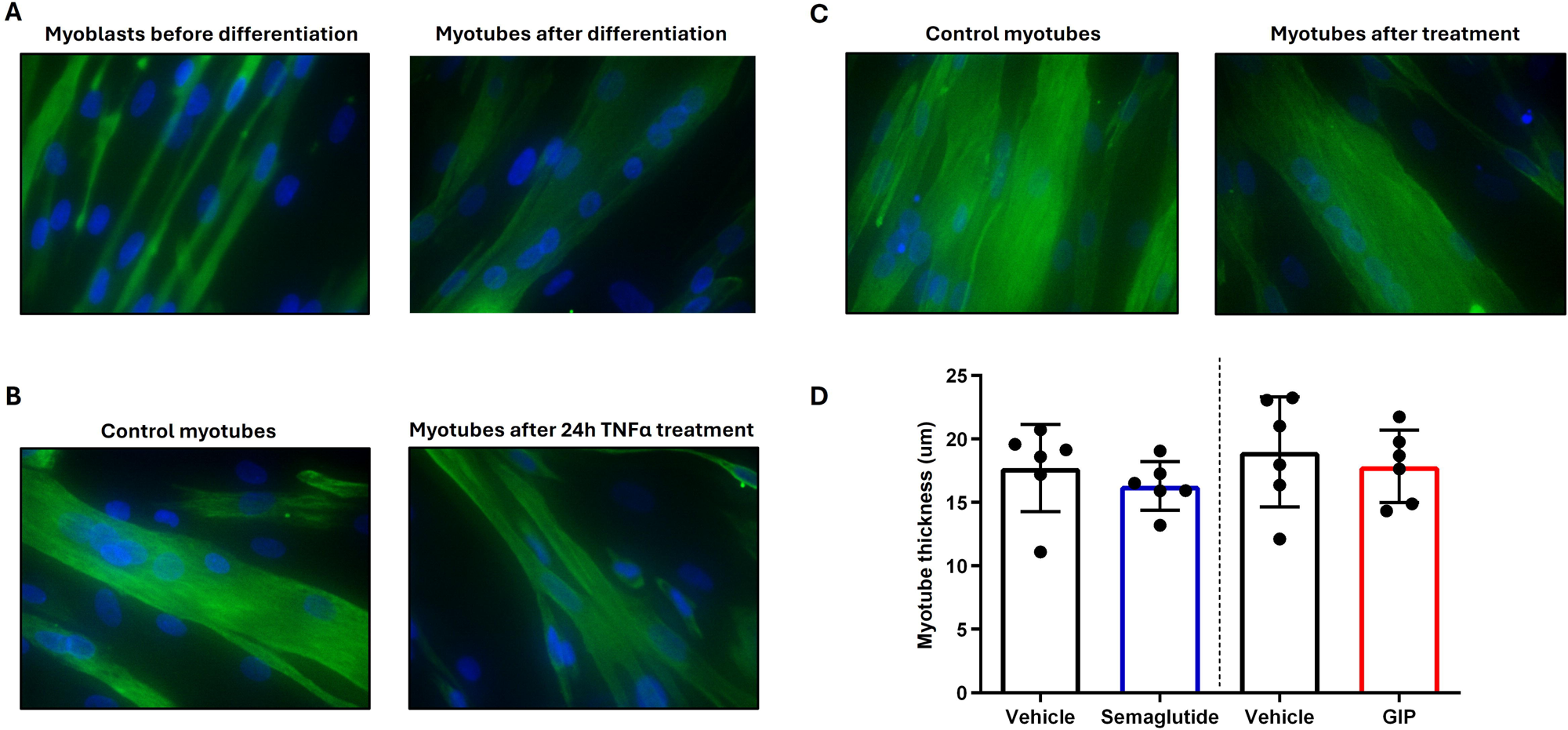
Incretins do not directly cause myotube atrophy. (A) Representative desmin/DAPI immunofluorescence images of human primary myoblasts before and after differentiation. Undifferentiated myoblasts appear as long, thin, mononucleated cells; differentiated myotubes are thicker and multinucleated. (B) Representative desmin/DAPI images of differentiated myotubes treated with or without TNFα for 24 hours. TNFα treatment induced visible myotube thinning while preserving multinucleation, confirming TNFα as a positive control for atrophy induction. (C) Representative desmin/DAPI images of control myotubes and myotubes following 24 hours of incretin treatment. (D) Myotube thickness (µm) following 24 hours of treatment with semaglutide vs vehicle (DMSO; left) or GIP vs vehicle (H_₂_O; right).

To assess whether direct incretin exposure influences myotube morphology, myotube thickness was quantified after 24 hours of semaglutide or GIP treatment. No visible morphological differences were observed between treatment and vehicle control conditions (Figure 4C), and quantitative analysis confirmed no significant effect of either semaglutide (p = 0.2824) or GIP (p = 0.5507) on myotube diameter compared with their respective vehicle controls (Figure 4D). These findings indicate that direct activation of incretin pathways does not induce myotube atrophy over 24 hours.

### Direct incretin treatment does not influence myogenesis

To assess whether incretins influence myogenic differentiation, myoblasts were treated with semaglutide or GIP every 48 hours throughout the 8-day differentiation protocol and myogenic marker expression was assessed by qPCR at day 2 and day 4. Expression of MyoD and Myf5 was not significantly altered by either treatment at either timepoint (Figure 5A). MyoG expression was not significantly affected by semaglutide at either day 2 or day 4 (Figure 5A). However, GIP treatment produced a statistically significant reduction in MyoG expression at day 4 compared with vehicle control (p = 0.0114), with a mean fold change of 0.77 relative to vehicle, representing a modest ∼23% reduction (Figure 5A). Given that this was not accompanied by changes in MyoD or Myf5, the functional significance of this observation is uncertain.

**Figure 5.**
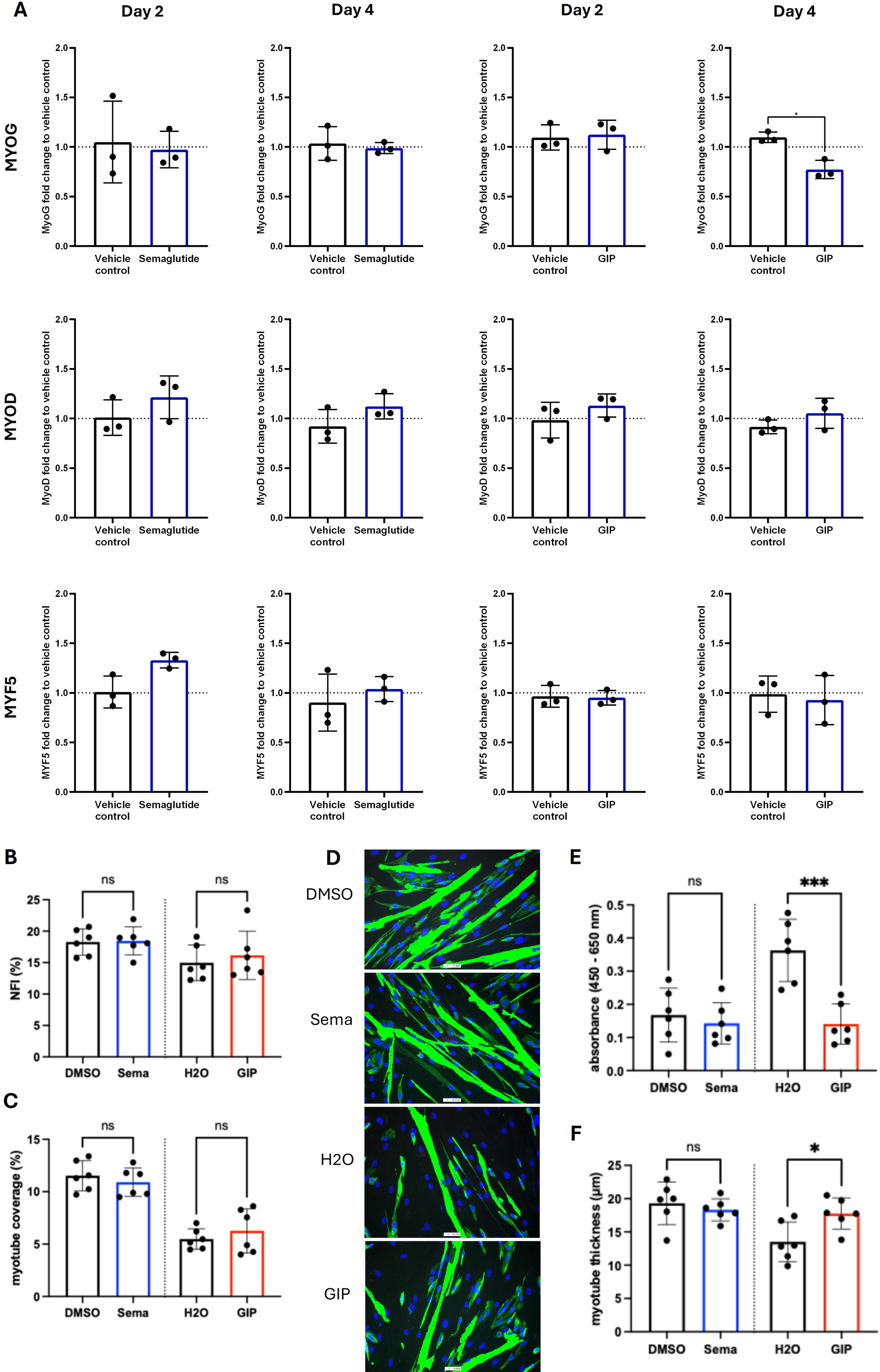
Incretins do not directly influence myogenesis. Data shown are from n = 3 biological replicates (PCR) or n = 6 biological replicates (all other assays). Myotubes were treated with semaglutide or GIP (with appropriate vehicle controls) every 2 days during the 8-day differentiation protocol. All comparisons by Wilcoxon signed-rank test; ns = not significant. (A) MyoG, MyoD, and Myf5 mRNA expression (fold change to vehicle control) at Day 2 and Day 4 of differentiation, for semaglutide vs vehicle control (left, blue) and GIP vs vehicle control (right, red). (B) Nuclear fusion index (NFI, %) following 8 days of differentiation with semaglutide vs DMSO (left) and GIP vs H_₂_O (right). Representative desmin/DAPI immunofluorescence images are shown for each condition (DMSO, semaglutide, H_₂_O, GIP). (C) Myotube coverage (%) following 8 days of differentiation. (D) Representative images used for lactate dehydrogenase (LDH) quantification, following 8 days of differentiation.| (E) LDH release (absorbance 450–650 nm) following 8 days of differentiation, shown for all four conditions. LDH release was consistent between semaglutide, DMSO, and GIP, but significantly elevated in the H_₂_O vehicle control condition relative to all three (vs DMSO, p = 0.0014; vs semaglutide, p = 0.0004; vs GIP, p = 0.0003), consistent with a hypotonic vehicle effect rather than a treatment effect. (F) Myotube thickness (µm) following 8 days of differentiation with semaglutide vs DMSO (left) and GIP vs H_₂_O (right). Semaglutide p = 0.7708; GIP p = 0.0205.

To assess the functional consequences of incretin treatment on differentiation outcomes, nuclei fusion index (NFI), myotube coverage, myotube thickness, and LDH release were quantified following 8 days of treatment. NFI and myotube coverage were not significantly affected by either semaglutide or GIP (NFI; semaglutide p = 0.9920, GIP p = 0.7293; Figure 5B; myotube coverage; semaglutide p = 0.7426, GIP p = 0.6236; Figure 5C). LDH release, a marker of cell damage, was consistent across semaglutide, DMSO (semaglutide vehicle control) and GIP treatments, but was significantly elevated by H2O (GIP vehicle control) treatment (vs DMSO, p = 0.0014; vs semaglutide, p = 0.0004; vs GIP, p = 0.0003; representative images shown in Figure 5D, absorbance values plotted in Figure 5E). We propose that hypotonic effects of the H2O vehicle control condition explain the reduction in myotube thickness. Myotube thickness was not significantly altered by semaglutide (p = 0.7708) but was significantly increased by GIP treatment, relative to vehicle control (p = 0.0205; Figure 5F).

## Discussion

The present study investigated whether GLP-1 and GIP receptor agonists exert direct pharmacological effects on primary human skeletal muscle, motivated by clinical observations that incretin-based therapies produce substantial reductions in lean body mass alongside fat mass. Our principal finding is that direct treatment with semaglutide (a GLP-1R agonist) or GIP peptide (a GIPR agonist) does not alter atrophic signalling, inflammatory gene expression, mitochondrial bioenergetics, myotube morphology, or myogenic differentiation in primary human myotubes derived from older adults. Semaglutide modestly reduced glycolytic ATP production and total ATP production rate, accompanied by reduced lactate secretion, though this was not of a magnitude consistent with clinically meaningful muscle impairment. To our knowledge these data provide the first experimental evidence that lean mass loss associated with GLP-1 and GIP/GLP-1 receptor agonist treatment in clinical trials is not driven by direct pharmacological action on skeletal muscle tissue.

At the level of nominal statistical thresholds (p < 0.05, log_₂_FC > 0.58), a small number of proteins warrant discussion. Following GIP treatment, three proteins were nominally upregulated: UBE2B (log_₂_FC = 0.65, p = 0.009), S100A12 (log_₂_FC = 0.87, p = 0.036), and CRNN (log_₂_FC = 1.05, p = 0.042). UBE2B is a ubiquitin-conjugating enzyme involved in proteasomal protein degradation and the DNA damage response; its modest upregulation could be consistent with proteostatic stress, though whether this is receptor-mediated or a non-specific effect of GIP cannot be determined from the current data. S100A12 is a calcium-binding protein of the S100 family with roles in inflammatory signalling and monocyte/macrophage recruitment; its elevation in conditioned media may reflect a secretory stress response, though it is not a canonical myokine. CRNN (cornulin) is an epithelial stress response protein with no established role in skeletal muscle. Following semaglutide treatment, only CD79B (log_₂_FC = 0.92, p = 0.014) was differentially expressed. CD79B is a component of the B-cell antigen receptor complex and has no known physiological role in skeletal muscle.

Several additional proteins showed nominal p < 0.05 without meeting the fold-change threshold and are worth highlighting due to their potential linkage to muscle mass and function. In the GIP comparison, ARG1 (log_₂_FC = 0.55, p = 0.011), arginase-1, regulates L-arginine metabolism and has been linked to inflammatory resolution in the context of macrophage-driven muscle regeneration (21); and DAPK2 (log_₂_FC = 0.42, p = 0.047), a death-associated protein kinase involved in autophagy regulation (22). For semaglutide, SAV1 was significantly increased (log_₂_FC = 0.48, p = 0.005), which is a scaffold protein in the Hippo signalling pathway that has been linked to muscle mass regulation and myogenesis (23, 24). Furthermore, there was a marginal but significant increase in FKBP5 (log_₂_FC = 0.43, p = 0.025), a glucocorticoid receptor co-chaperone implicated in stress response and metabolic regulation (25), and GPD1 (log_₂_FC = 0.36, p = 0.014), a glycerol-3- phosphate dehydrogenase involved in lipid metabolism and redox balance (26). Whilst individually none of these protein changes met our thresholds of differential expression, their alteration and potential linkage to pathways that mediate muscle mass and functional might warrant targeted follow-up in a more powered study. Critically, none of the six proteins with established relevance to muscle atrophy, hypertrophy, and inflammation (namely FOXO1, MSTN, IGF1R, TGFB1, FOXO3, and IL6) showed statistically significant changes in either comparison, with all nominal p-values exceeding 0.05. The closest to nominal significance were IGF1R (semaglutide: p = 0.057) and FOXO3 (semaglutide: p = 0.060), both showing modest increases in the semaglutide condition that did not reach the conventional threshold. Taken together, the Olink data support the conclusion that direct incretin exposure does not meaningfully remodel the muscle secretory proteome and, in particular, does not engage canonical pathways governing muscle mass or inflammation.

A key preliminary finding of this study was the differential expression of incretin receptors in human primary myotubes. GIPR mRNA was consistently detected across all six donors, whilst GLP-1R mRNA was undetectable by RT-PCR. This is consistent with published transcriptomic datasets showing that GIPR is expressed at low but detectable levels in human skeletal muscle, whilst GLP-1R expression is generally below reliable detection thresholds in muscle tissue (15, 17, 27). The use of LUXendin645, a validated, cell-impermeant fluorescent GLP-1R antagonist probe (20), allowed us to interrogate surface GLP-1R protein independently of mRNA detection. Flow cytometric analysis revealed a significant increase in LUXendin645 mean fluorescence intensity relative to unstained controls, with 51–66% of myoblasts classified as probe-positive, suggesting that a proportion of cells express low levels of surface GLP-1R protein. This discordance between mRNA and protein is not unusual for low-abundance receptors and has been noted in other cell types (28). Nonetheless, the very low mRNA levels observed suggest that GLP-1R expression in human myotubes is likely to be functionally limited, which is consistent with the absence of significant biological responses to semaglutide observed across our functional assays.

In the context of lean mass loss with incretin agonists, our data support the view that the mechanism is indirect rather than a direct pharmacological effect on muscle. In the major clinical trials, the proportion of lean mass lost relative to total body-weight (∼25%) is broadly consistent with that observed in diet-induced weight loss studies (29), suggesting that incretin receptor agonism does not disproportionately drive muscle catabolism beyond what would be expected from the caloric deficit alone. The reduction in mechanical loading that accompanies lower body weight may also contribute to lean mass loss through the induction of disuse-atrophy catabolic mechanisms (30, 31), particularly in older or less active individuals. Our finding that neither semaglutide nor GIP treatment altered the expression of canonical atrophy-related genes (MAFBX, MURF1, FOXO3) or induced morphological changes in myotube diameter is consistent with this interpretation and argues against a direct atrophic stimulus from incretin receptor activation.

We did observe a modest but statistically significant reduction in glycolytic ATP production rate and total ATP production rate in myotubes treated with semaglutide, which was supported by a significant reduction in lactate secretion. These findings are mechanistically coherent, suppression of glycolytic flux would be expected to reduce pyruvate availability and, consequently lactate production via lactate dehydrogenase. GLP-1R signalling is known to modulate glycolytic activity in pancreatic beta cells (32)and adipocytes (33), where cAMP-mediated signalling can influence glucose transporter trafficking and glycolytic enzyme activity (32). Whether a similar mechanism operates in skeletal muscle where the GLP-1R is expressed at low levels remains unclear. Critically, these glycolytic changes were not accompanied by any perturbation of mitochondrial ATP production, oxygen consumption rate, or coupling efficiency, and the overall magnitude was small. The effect is therefore unlikely to carry clinically meaningful consequences for skeletal muscle energetics, though whether sustained in vivo dosing over weeks to months might amplify such adaptations cannot be excluded.

The myogenesis data offered an intriguing contrast between the two incretin treatments. Whilst neither semaglutide nor GIP altered nuclei fusion index, myotube coverage, or the expression of the early myogenic transcription factors MyoD and Myf5 during differentiation, GIP treatment produced a statistically significant reduction in myogenin (MyoG) expression at Day 4 of differentiation. Myogenin is a terminal differentiation factor required for myoblast fusion and myotube maturation, and its reduced expression might, in principle, be expected to impair differentiation. However, this was not borne out functionally: nuclei fusion index and myotube coverage at Day 8 were unaffected. This dissociation suggests either that MyoG expression was transiently suppressed without reaching a threshold sufficient to alter differentiation outcomes, or that compensatory mechanisms maintained normal differentiation kinetics. Notably, after 8 days of differentiation, GIP treatment significantly increased myotube thickness whilst simultaneously reducing LDH release relative to vehicle control. This combination of findings is more consistent with a modest pro-hypertrophic or cytoprotective effect of GIPR activation during myogenesis, resulting in larger myotubes with less evidence of cell damage. Taken together, these findings do not indicate a detrimental pro-atrophic or anti-myogenic effect of GIP peptide or semaglutide on skeletal muscle under the conditions studied.

The observation that GIP treatment significantly increased IL-6 secretion at 2 hours by ELISA, a finding not reflected in the 4-hour gene expression data or the Olink secretome analysis at 24 hours, deserves comment. IL-6 is a pleiotropic cytokine with both pro- and anti-inflammatory roles in skeletal muscle. When secreted acutely by contracting muscle, it acts as a myokine with metabolic and anti-inflammatory signalling functions; conversely, chronically elevated systemic IL-6 contributes to muscle catabolism in inflammatory conditions (34).

Whether GIPR activation on myotubes can directly stimulate acute IL-6 release is a novel finding that merits replication and mechanistic follow-up. The transient nature of the observed response is more consistent with a brief, auto-regulated signal rather than a sustained pro-inflammatory response. A receptor-mediated mechanism involving cAMP-dependent transcriptional activation is biologically plausible given the known downstream signalling of GIPR (35), but remains to be directly demonstrated in skeletal muscle.

This study has several limitations that should be considered when interpreting the findings. First, the myotubes used were derived from older adults undergoing elective joint replacement surgery for end-stage osteoarthritis, a condition associated with chronic systemic inflammation, altered muscle metabolism, and age-related skeletal muscle changes. Although myotubes derived from OA patients are a well-established and widely used model system (36–41), the extent to which they fully recapitulate muscle biology in younger or healthier individuals is uncertain. Second, the in vitro treatment employed here inevitably differs somewhat from the pharmacokinetic and pharmacodynamic profile of incretin agonists administered chronically in vivo. The concentrations used (semaglutide 10 nM; GIP 5.02 μM) were selected to reflect circulating therapeutic ranges and those used in prior experimental studies, but cells were exposed for 24 hours in isolation, without the complex endocrine and paracrine environment of intact muscle tissue. In vivo, muscle is embedded within a milieu of adipokines, hormones, neural inputs, and mechanical stimuli that may modulate its sensitivity to incretin receptor signalling. Third, the study was powered to detect substantial effects and, with n = 6 donors for most assays, may have been underpowered to detect modest but reproducible changes, particularly in the secretome analysis with over 1,000 proteins tested simultaneously. Fourth, whilst the 24-hour and 8-day treatment windows used here provide snapshots of acute and differentiation-phase responses respectively, the effects of long-term incretin exposure on muscle cannot be inferred from these data.

## Conclusion

In conclusion, this study provides evidence that GLP-1 and GIP receptor agonism does not directly drive skeletal muscle atrophy, inflammatory signalling dysregulation, or myogenic impairment in primary human myotubes. The reduction in lean mass observed clinically with incretin-based therapies is therefore more consistent with an indirect mechanism, most likely secondary to the caloric deficit and reduced mechanical load accompanying weight loss, rather than a direct pharmacological effect on muscle tissue. Whilst a modest suppression of glycolytic flux with semaglutide and a transient IL-6 response with GIP were observed, neither finding was of a magnitude or character that would indicate a clinically concerning direct muscle effect. These findings are of importance as the use of incretin agonists expands to older and frailer populations at risk of sarcopenia, providing some reassurance that these agents do not directly accelerate muscle wasting. Future work should examine whether long-duration or repeated incretin exposure in more complex three-dimensional muscle models or ex vivo preparations alters muscle biology in ways not captured by the current experimental system.

## List of abbreviations

2NBDG: 2-(N-(7-nitrobenz-2-oxa-1,3-diazol-4-yl)amino)-2-deoxyglucose
ATP: adenosine triphosphate
BH: Benjamini–Hochberg
BMI: body mass index
BSA: bovine serum albumin
CD79B: CD79b molecule
CRNN: cornulin
CT/ΔCT: cycle threshold/delta cycle threshold
DAPI: 4′,6-diamidino-2-phenylindole
DMSO: dimethyl sulfoxide
DPP-4: dipeptidyl peptidase-4
FOXO1/FOXO3: forkhead box O1/O3
GIP: glucose-dependent insulinotropic polypeptide
GIPR: GIP receptor
GLP-1: glucagon-like peptide-1
GLP-1R: GLP-1 receptor
GlycoPER: glycolytic proton efflux rate
IGF1R: insulin-like growth factor 1 receptor
IL-6/IL6: interleukin-6
J_ATP-GLYC: glycolytic ATP production rate
J_ATP-MITO: mitochondrial ATP production rate
LDH: lactate dehydrogenase
log_₂_FC: log_₂_ fold change
MAFBX: muscle atrophy F-box (atrogin-1)
MFI: mean fluorescence intensity
MSTN: myostatin
MURF1: muscle RING-finger protein 1
Myf5: myogenic factor 5
MyoD: myoblast determination protein 1
MyoG: myogenin
NFI: nuclear fusion index
NPX: normalised protein expression
OCR: oxygen consumption rate
PBS: phosphate-buffered saline
PCA: principal component analysis
PCR/RT-qPCR: polymerase chain reaction/reverse transcription quantitative PCR
S100A12: S100 calcium-binding protein A12
SD: standard deviation
SEM: standard error of the mean
TGFB/TGFB1: transforming growth factor beta 1
TNFα: tumour necrosis factor alpha
UBE2B: ubiquitin-conjugating enzyme E2 B

## Acknowledgments

The authors thank all patients who donated tissue samples for this research, and the clinical and surgical teams at the Royal Orthopaedic Hospital (Birmingham) and Russell’s Hall Hospital (Dudley) for their assistance with sample collection. This research is funded by the National Institute for Health and Care Research (NIHR) Birmingham Biomedical Research Centre (BRC). The views expressed are those of the author(s) and not necessarily those of the NIHR or the Department of Health and Social Care (Award reference NIHR-INF-5081). The authors would also like to acknowledge Dr Jon Hazeldine for guidance and providing the 2NBDG.

---

**Supplementary figure 1.** Quality control of Olink Reveal proteomic data. **(A)** Kernel density plots showing the distribution of normalised protein expression (NPX) values for each sample, colour-coded by treatment group. Overlapping distributions across samples confirm consistent data quality. **(B)** Box-and-whisker plots showing the distribution of NPX values for each individual sample (labelled by sample ID), colour-coded by treatment group.

