## Supplemental Table 1 for "GLP-1 and GIP receptor agonism does not directly drive skeletal muscle atrophy or impair myogenesis in primary human myotubes"

**Supplementary**

Table S1. TaqMan probe assay IDs and SYBR green gene-specific primer sequences used for RT-qPCR.

| Gene | Supplier | Assay ID/sequence |
| --- | --- | --- |
| GAPDH | Thermofisher | **Hs02786624_g1** |
| GIPR | Thermofisher | **Hs00609201_g1** |
| GLP-1R | Thermofisher | **Hs00157705_m1** |
| MAFbx | Thermofisher | **Hs01041408_m1** |
| MuRF1/Trim63 | Thermofisher | **Hs00822397_m1** |
| Foxo3 | Thermofisher | **Hs00818121_m1** |
| IL6 | Thermofisher | **Hs00174131_m1** |
| TGFB/Tgfb1 | Thermofisher | **Hs00998133_m1** |
| MYOG | Primerdesign | Forward: 5'-GCCCTGATGCTAGGAAGCC-3' Reverse: 5'-CTGAATGAGGGCGTCCAGTC-3' |
| MyoD | Primerdesign | Forward: 5'-CGCCTGAGCAAAGTAAATGAG-3' Reverse: 5'-GCCCTCGATATAGCGGATG-3' |
| MYF5 | Primerdesign | Forward: 5'-CACCTCCAACTGCTCTGATG-3' Reverse: 5'-TAAGGAGTTTTATCTGTGGCATATAC-3' |
| GAPDH | Primerdesign | Proprietary |
